# Neural mechanisms underlying reward processing and social cognition: a replication study with a Japanese sample

**DOI:** 10.1101/2025.07.03.662916

**Authors:** Kenta Ohnishi, Michiyo Sugawara, Yoko Mano, Shinsuke Suzuki

## Abstract

Neural functions underlying reward processing and social cognition play a critical role in everyday decision-making. Given that these processes may be shaped by cultural factors, it is essential to examine their cross-cultural generalizability. In this study, we used functional MRI to scan native Japanese speakers as they performed two well-established experimental paradigms: the Monetary Incentive Delay (MID) task for reward processing and the Theory of Mind (ToM) task for social cognition. We successfully replicated previous findings. Specifically, in the MID task, reward expectation and reward outcome were associated with neural activity in the ventral striatum and ventromedial prefrontal cortex. In the ToM task, social cognition was linked to activation in the temporoparietal junction. Notably, the posterior cingulate cortex was engaged in both tasks, suggesting its integrative role across cognitive domains. Together, these results replicate and extend earlier work, supporting the cross-cultural generalizability of the neural mechanisms underlying reward and social cognition, and further validating our fMRI protocol for future research.

## Introduction

A new interdisciplinary field known as neuroeconomics, which integrates neuroscience and economics, emerged in the early 2000s (Fehr and Camerer 2007; Glimcher 2013). A series of studies in this field has elucidated the neural mechanisms underlying key cognitive and emotional processes that have been extensively examined in social sciences, particularly focusing on themes such as reward processing and social cognition. These studies have implicated a brain network including the ventral striatum and ventromedial prefrontal cortex (vmPFC) in reward processing (Bartra, McGuire, and Kable 2013; Clithero and Rangel 2014; Suzuki 2022), and another network including the temporoparietal junction (TPJ) in social cognition (Frith and Frith 2012; Van Overwalle 2009; Suzuki and O’Doherty 2020).

Our economic decisions are guided by expected rewards. For instance, investment behavior relies on expectations about whether the value of an asset will rise or fall. The expectations can subsequently be revised in response to the outcomes of realized rewards. Accumulating evidence from human neuroimaging studies employing the Monetary Incentive Delay (MID) task has demonstrated that the ventral striatum conveys information about expected rewards, while the vmPFC encodes reward outcomes (Knutson et al. 2000; Oldham et al. 2018; Jauhar et al. 2021). Consistent with these findings, meta-analyses have repeatedly identified these two regions as central components of the brain’s reward processing network (Bartra, McGuire, and Kable 2013; Clithero and Rangel 2014). The MID task is also widely used to investigate individual differences in neural reward sensitivity and their relationships with various mental disorders and personality traits (Ng, Alloy, and Smith 2019; Matyjek, Bayer, and Dziobek 2023).

Economic decisions often involve social contexts. Therefore, the ability to infer others’ intentions, beliefs, and mental states is crucial for making appropriate decisions. For example, in stock trading, it is important to consider what other investors believe about market conditions, while in business decision-making, anticipating competitors’ next actions is essential. Brain regions associated with such social inference—also referred to as social cognition—have been localized using the Theory of Mind (ToM) task (Dodell-Feder et al. 2011; Ogawa, Yokoyama, and Kameda 2017). A large body of research using this task has shown that social cognition recruits multiple brain regions, including the TPJ, dmPFC, temporal pole, and posterior cingulate cortex (PCC) (Schurz et al. 2014). The ToM task is also commonly used in psychiatric and developmental research to explore individual differences in the neural basis of social cognition (Dodell-Feder, DeLisi, and Hooker 2014; Mahy, Moses, and Pfeifer 2014).

While reward processing and social cognition both play pivotal roles in economic decision-making, less is known about whether there is a brain region commonly involved in both functions. There is still a lack of research in which both the ToM task and the MID task are administered to the same participants (Erk et al. 2017). Examining both functions within the same participants may offer a more unified account of the neural architecture supporting economic decision-making.

In this study, we aimed to replicate previous findings from the MID and ToM tasks in a Japanese sample, using a newly installed MRI scanner at Hitotsubashi University in Japan. Replication is a critical step in advancing scientific knowledge, as it allows researchers to evaluate the generalizability of original findings across different settings and populations (Munafò et al. 2017). This is particularly important for constructs such as social cognition, which may vary depending on cultural and linguistic context (Kitayama and Uskul 2011; Park and Huang 2010). Given that cultural factors shape both social cognition and valuation processes, it is essential to examine the replicability of the neural correlates of social cognition across diverse populations (Han and Northoff 2008). Furthermore, to date, no study has investigated neural activity during both the MID and ToM tasks within the same participants using a Japanese sample.

In our experiment, the same group of participants completed both the MID and ToM tasks while undergoing functional MRI scanning. In the MID task, we observed neural activity associated with reward expectation in the bilateral ventral striatum, and activity related to reward outcomes in the ventromedial prefrontal cortex (vmPFC). In the ToM task, we identified neural activation associated with social cognition in the temporoparietal junction (TPJ). Notably, we also found significant activation in the posterior cingulate cortex (PCC) in response to both reward outcomes and social cognition, suggesting a potential integrative role for this region in economic decision-making.

## Methods

This study was approved by the Ethics Committee of Hitotsubashi University (ID: 2024C051).

### Participants

Fourteen participants (thirteen males; age = 25.64 ± 6.74 years, mean ± SD) completed both the Monetary Incentive Delay (MID) task and the Theory of Mind (ToM) task from April 30 to May 16, 2025. All participants were native Japanese speakers, and pre-screened to exclude individuals with a history of neurological or psychiatric illness. Informed written consent was obtained from all participants. They received monetary rewards based on their performance in the MID task (see below), in addition to a participation fee of 4,000 Japanese Yen (JPY).

### Experimental tasks

Participants performed the MID and ToM tasks in the fixed order.

#### MID task

In this task (Knutson et al. 2000), participants were required to press a button as quickly as possible when a target shape appeared (Fig. 1a). Each trial began with an inter-trial interval (ITI) lasting 2–12 seconds, followed by a cue indicating the amount of monetary reward available upon a successful response (Cue phase, 1 second). After a short inter-stimulus interval (ISI) of 2–2.5 seconds, a square target shape was presented (Target phase, approximately 0.25 seconds). If the participant pressed the button within the time limit (i.e., while the target was on screen), the trial was considered successful, and the participant received the indicated reward. If the response was too late, no reward was given. The time limit (i.e., target presentation duration) was individually adjusted to maintain a success rate of approximately 0.66.

**Figure 1:**
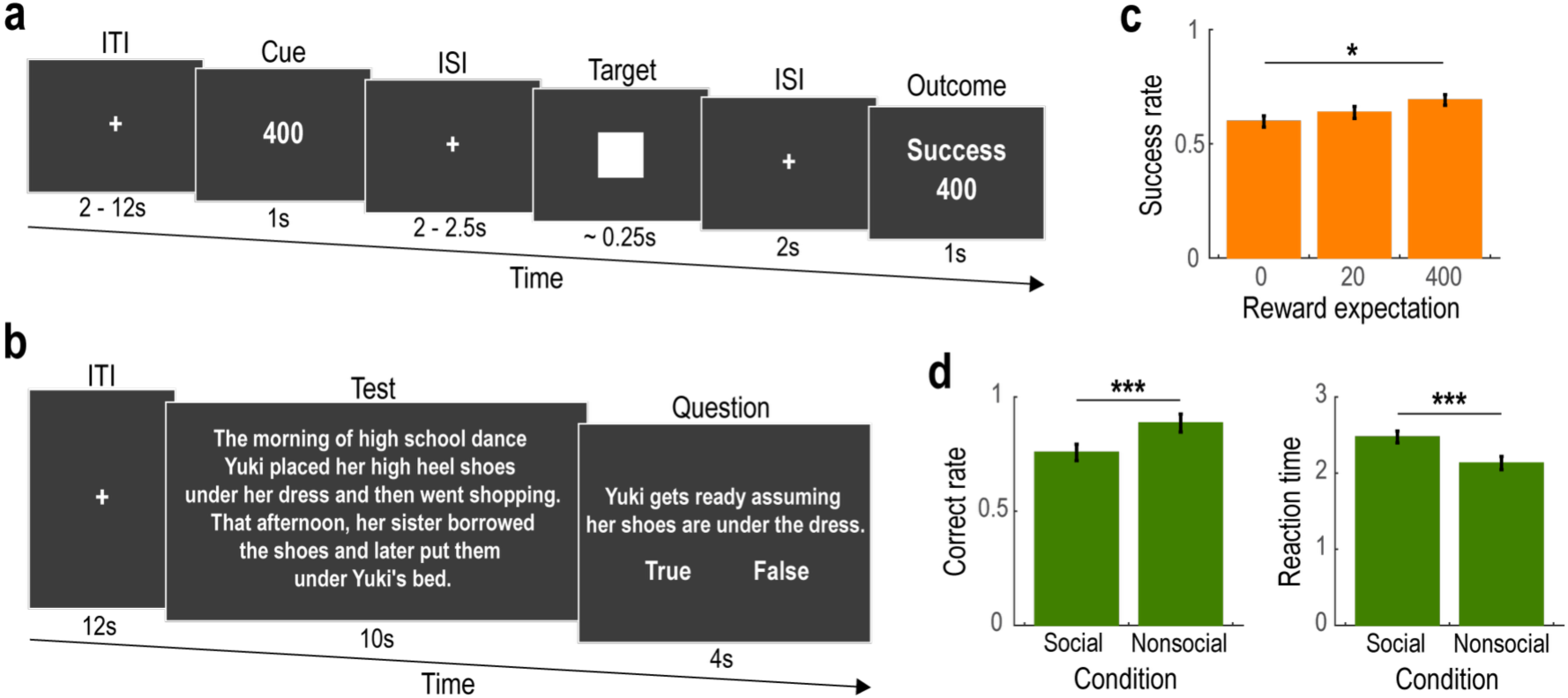
Experimental task and behavior. (a) Illustration of the Monetary Incentive Delay (MID) task. On each trial, participants were instructed to press a button as quickly as possible when a target rectangle appeared. If the button was pressed within the time limit (i.e., while the target was on screen), the trial outcome was considered successful, and the participant received the monetary reward indicated in the Cue phase. ITI, inter-trial-interval; ISI, inter-stimulus-interval. (b) Illustration of the Theory of Mind (ToM) task. On each trial, participants read a sentence stimulus presented on a computer screen and responded by pressing a button to indicate whether the content was “True” or “False.” The sentence set included 10 descriptions about social situations (Social condition) and 10 about nonsocial situations (Nonsocial condition). (c) Success rate in the MID task (Mean ± SEM across participants). Success rates are plotted as a function of reward expectation (i.e., the monetary amount available upon a successful outcome). \**p* < 0.05. (d) Correct rate and reaction time in the ToM task (Mean ± SEM across participants). The data are shown separately for the Social and Nonsocial conditions. *Left*, correct rate. *Right*, reaction time. \*\*\**p* < 0.001.

The initial duration was set to 0.25 seconds and was adaptively updated based on performance: if the correct response rate up to the current trial exceeded 66%, the time limit was reduced by 0.025 seconds in the next trial; if it fell below 66%, the time limit was increased by 0.025 seconds. Finally, participants received feedback indicating the success or failure of their response (Outcome phase, 1 second).

Participants completed two runs of the task, each consisting of 30 trials. The reward amount varied across trials (0 JPY, 20 JPY, or 400 JPY), and the order of reward conditions was randomized across participants.

#### ToM task

In this task (Dodell-Feder et al. 2011; Ogawa, Yokoyama, and Kameda 2017), participants read sentences presented on a computer screen and responded by pressing a button to indicate whether the content was “True” or “False” (Fig. 1b). Each trial began with an inter-trial interval (ITI) of 12 seconds, followed by the presentation of a sentence stimulus (Text phase, 10 seconds). Participants were then prompted to judge the truthfulness of the statement (Question phase, 4 seconds). No feedback was provided regarding their responses. The sentence set consisted of 20 stimuli: 10 described social situations (Social condition: e.g., “*The morning of high school dance Yuki placed her high heel shoes under her dress and then went shopping. That afternoon, her sister borrowed the shoes and later put them under Yuki’s bed. Yuki gets ready assuming her shoes are under the dress.*”) and 10 described nonsocial situations (Nonsocial condition: e.g., “*Part of the garden is supposed to be reserved for the roses; it’s labeled accordingly. Recently the garden has run wild, and dandelions have taken over the entire flower bed. According to the label, these flowers are roses.*”). The sentences were presented in random order across participants. See the article (Ogawa, Yokoyama, and Kameda 2017) for the full list of sentence stimuli.

### Reward payment

Participants received a participation fee of 4,000 JPY. In addition, they were awarded a performance-based bonus from the MID task. Specifically, at the end of the experiment, one trial was randomly selected, and the monetary reward from that trial was implemented as an additional payment.

### Behavioral data analysis

#### MID task

To examine the effect of reward expectation on the success rate, we conducted a generalized linear mixed-effects model (GLMM) analysis:

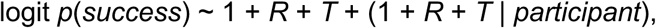

where *R* denotes the amount of monetary reward presented during the Cue phase, and *T* represents the trial number (i.e., 1 for the first trial, 2 for the second trial, and so on). We included the trial number, *T*, to assess the effect of experience (learning).

*ToM* task. We conducted GLMM analyses to examine the effects of experimental condition on correct rate and reaction time. The models were specified as follows:

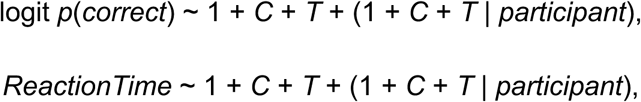

where *C* indicates the condition (1 for Social, 0 for Nonsocial), and *T* again represents the trial number.

### fMRI data acquisition

We collected fMRI images using a 3T Siemens MAGNETOM Prisma scanner located at the HIAS Brain Research Center (Hitotsubashi University, Tokyo, Japan) with a 32-channel radio frequency coil. The BOLD signal was measured using a T2*-weighted gradient-echo echo-planar imaging (EPI) sequence with the following parameters: TR = 800 ms, TE = 34.4 ms, flip angle = 52°, voxel size = 2.4 × 2.4 × 2.4 mm, matrix size = 86 × 86, 60 slices aligned to the AC–PC plane, multiband acceleration factor = 6. Before the functional runs, high-resolution (0.9 mm^3^) anatomical images were acquired using a standard MPRAGE pulse sequence (TR = 2,300 ms, TE = 2.32 ms, FA = 8°).

### fMRI data preprocessing

Functional MRI data were preprocessed using standard procedures implemented in SPM12. For each participant, images were realigned to the first volume to correct for head motion, spatially normalized to the MNI template, and temporally filtered using a high-pass filter with a cutoff of 128 seconds. Spatial smoothing was applied using an 8-mm full-width at half-maximum (FWHM) Gaussian kernel. In addition, geometric distortions in the fMRI images were corrected using the SynBOLD-DisCo method after the realignment (Yu et al. 2023).

### fMRI data analysis

To identify the neural correlates of reward processing and social cognition, we performed standard General Linear Model (GLM) analyses.

#### GLM for the MID task

We defined a separate GLM for each participant. The design matrix included the following regressors: (1) a boxcar function representing the Cue phase, (2) a boxcar function for the Target phase, and (3) two boxcar functions for the Outcome phase, modeled separately for success and failure trials. In addition, a parametric modulator was included for the Cue phase boxcar function, encoding the amount of monetary reward that could be obtained upon a successful outcome. All regressors were convolved with a canonical hemodynamic response function (HRF).

Six motion-correction parameters were included as regressors of no interest to account for head movement artifacts. We defined two contrasts of interest: reward expectation (i.e., the parametric modulator during the Cue phase) and reward outcome (i.e., the contrast between success and failure trials during the Outcome phase). For each participant, contrast images were estimated at every voxel across the whole brain and then submitted to a random-effects group-level analysis.

#### GLM for the ToM task

The design matrix included two regressors: boxcar functions covering the Text and Question phases for the Social and Nonsocial conditions, respectively. Following previous studies (Ogawa, Yokoyama, and Kameda 2017), we modeled neural activity across the Text and Question phases as a single regressor for each condition. Six motion-correction parameters were included as regressors of no interest. The primary contrast of interest for social cognition was the comparison between the Social and Nonsocial conditions.

#### Whole-brain analysis

We set the significance threshold at *p* < 0.05, corrected for multiple comparisons at the cluster level across the whole brain, using a cluster-forming threshold of *p* < 0.001 (uncorrected) (Eklund, Nichols, and Knutsson 2016). The unthresholded activation maps are available at NeuroVault (https://neurovault.org/collections/21164/).

#### ROI analysis

Regions of interest (ROIs) were defined based on association maps from Neurosynth (https://neurosynth.org/) (Yarkoni et al. 2011). We used the terms “reward” and “social” to identify peak voxels associated with reward processing and social cognition, respectively. Spherical ROIs (10 mm radius) were then centered on these peak coordinates (reward-related ROIs: ventromedial prefrontal cortex, [x, y, z] = [2.0, 58.0, −8.0]; right ventral striatum, [12.0, 10.0, −8.0], and left ventral striatum, [−12.0, 10.0, −8.0]; and social cognition-related ROIs: right temporoparietal junction, [50.0, − 44.0, 10.0], left temporoparietal junction, [−56.0, −58.0, 20.0], right temporal pole, [52.0, 8.0, −32.0], left temporal pole, [−50.0, 16.0, −28.0], dorsomedial prefrontal cortex, [2.0, 56.0, 20.0], posterior cingulate cortex, [−2.0,-56.0,36.0]). Statistical significance was assessed at *p* < 0.05 (uncorrected), and results were also evaluated using Bonferroni correction for multiple comparisons across ROIs.

## Results

### Behavior

In the Monetary Incentive Delay (MID) task, participants successfully pressed the button within the time limit in approximately 64% of trials (success rate = 0.641 ± 0.010, mean ± SEM across participants). Furthermore, the success rate was significantly modulated by reward expectation—that is, the amount of money obtained upon a successful outcome (see Fig. 1c; effect of reward expectation: β = 0.089 x 10^-2^ ± 0.040 x 10^-2^, *t* = 2.188, and *p* = 0.029). There was no significant improvement in the success rate over time (effect of trial number: β = −0.066 x 10^-2^ ± 0.416 x 10^-2^, *t* = − 0.160, and *p* = 0.873).

In the Theory of Mind (ToM) task, participants gave the correct answer in approximately 82% of trials (correct rate = 0.821 ± 0.029). When comparing the Social and Nonsocial conditions, the correct rate was slightly higher in the Nonsocial condition (see Fig. 1d, left; effect of condition: β = −0.988 ± 0.416, *t* = −2.373, and *p* = 0.018). Reaction time was also shorter in the Nonsocial than in the Social condition (see Fig. 1d, right; effect of condition: β = 0.336 ± 0.066, *t* = 5.059, and *p* < 0.001).

Neither the correct rate nor reaction time was significantly improved over time (effect of trial number on correct rate: β = 0.055 ± 0.039, *t* = 1.394, and *p* = 0.164; effect of trial number on reaction time: β = 0.007 ± 0.006, *t* = −1.451, and *p* = 0.148).

### Neuroimaging: MID task

We first identified neural correlates of reward expectation (i.e., the parametric modulation by the amount of monetary reward presented at the Cue phase) and outcomes (i.e., the contrast between success and failure at the Outcome phase) in the MID task. Whole-brain analysis revealed that, at the time of cue presentation, reward expectation was encoded in a large cluster including the ventral striatum (Fig. 2a, p < 0.05, whole-brain corrected at the cluster level; see Table S1 for other activated regions). The BOLD signal within this cluster was significantly correlated with the amount of money obtainable upon a successful outcome during the Cue phase. Further analysis on independently defined regions of interest (ROIs) demonstrated that the left and right ventral striatum, but not the ventromedial prefrontal cortex (vmPFC), significantly represented reward expectation (Fig. 2b: *t* = 4.972, *p* = 0.006 in the left ventral striatum; *t* = 5.073, *p* = 0.002 in the right ventral striatum; *t* = 1.967, *p* = 0.070 in the vmPFC). Notably, the BOLD signal in the ventral striatum varied in a graded manner according to the magnitude of reward expectation (Fig. S1a).

**Figure 2:**
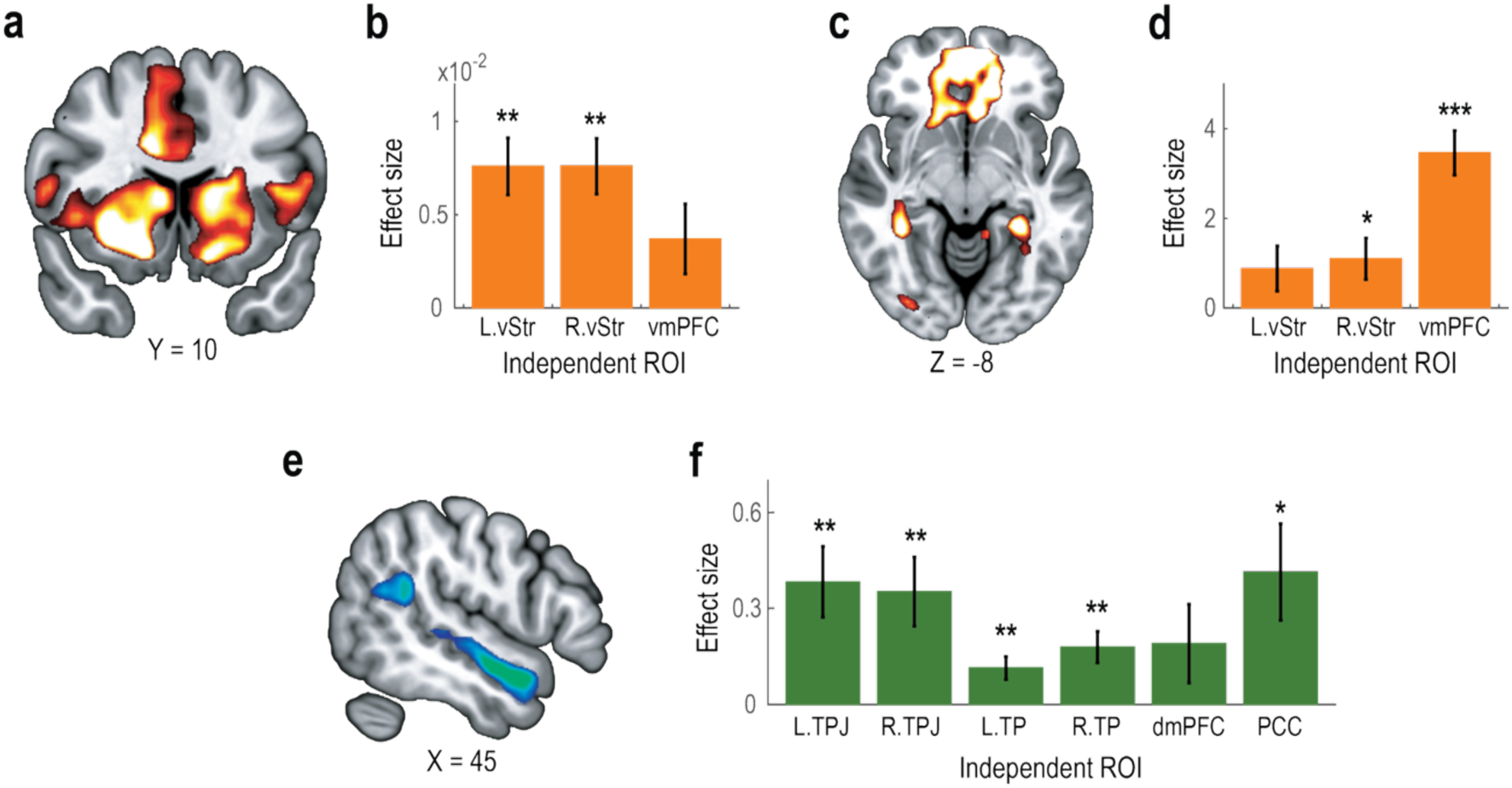
Neuroimaging results. (a) Whole-brain analysis of neural correlates of reward expectation in the MID task. Statistical parametric map showing neural activity parametrically modulated by the amount of monetary reward presented during the Cue phase. The activation map is thresholded at *p* < 0.001 (uncorrected) for display purposes. (b) Independent ROI analysis of reward expectation in the MID task. Bar plots show the effect sizes of reward expectation (Mean ± SEM across participants) in independently defined regions of interest. L.vStr, left ventral striatum; R.vStr, right ventral striatum; and vmPFC, ventromedial prefrontal cortex. \*\**p* < 0.01. (c) Whole-brain analysis of neural correlates of reward outcome in the MID task. Statistical map showing neural activity associated with the contrast between success and failure during the Outcome phase. The activation map is thresholded at *p* < 0.001 (uncorrected) for display purposes. (d) Independent ROI analysis of reward outcome in the MID task. Bar plots show the effect sizes of reward outcome (Mean ± SEM across participants). \**p* < 0.05 and \*\*\**p* < 0.001. (e) Whole-brain analysis of neural correlates of social cognition in the ToM task. Statistical map showing neural activity associated with the contrast between Social and Nonsocial conditions. The activation map is thresholded at *p* < 0.001 (uncorrected) for display purposes. (f) Independent ROI analysis of social cognition in the ToM task. Bar plots show effect sizes of social cognition (Mean ± SEM across participants). L.TPJ, left temporoparietal junction; R.TPJ, right temporoparietal junction; L.TP, left temporal pole; R.TP, right temporal pole; dmpFC, dorsomedial prefrontal cortex; and PCC, posterior cingulate cortex.

Whole-brain analysis of reward outcomes revealed that the vmPFC showed greater neural activity for successful compared to unsuccessful trials during the Outcome phase (Fig. 2c, p < 0.05, whole-brain corrected at the cluster level; see Table S1 for other activated regions). Independent ROI analyses further demonstrated that, in addition to the vmPFC, the right ventral striatum also represented reward outcomes (Fig. 2d: *t* = 6.947, *p* < 0.001 in the vmPFC; *t* = 2.363, *p* = 0.034 in the right ventral striatum; *t* = 1.740, *p* = 0.105 in the left ventral striatum). See Fig. S1b for separate visualizations of neural activation in success and failure trials.

### Neuroimaging: ToM task

In the ToM task, whole-brain analysis revealed that neural activity in the right temporoparietal junction (TPJ) and temporal poles (TP) was associated with social cognition (Fig. 2e, p < 0.05, whole-brain corrected at the cluster level; see Table S2 for other activated regions). BOLD signals in the TPJ and TP were significantly elevated in the Social condition compared to the Nonsocial condition. In addition to these areas, independent ROI analyses showed that the posterior cingulate cortex (PCC), but not the dorsomedial prefrontal cortex (dmPFC), exhibited differential activation between the Social and Nonsocial conditions (Fig. 2f: *t* = 3.466, *p* = 0.004 in the left TPJ; *t* = 3.267, *p* = 0.006 in the right TPJ; *t* = 3.161, *p* = 0.008 in the left TP; *t* = 3.643, *p* = 0.003 in the right TP; *t* = 1.547, *p* = 0.146 in the dmPFC; *t* = 2.743, *p* = 0.017 in the PCC). See Fig. S1c for neural responses in the Social and Nonsocial conditions displayed separately.

### Neuroimaging: shared representation between the MID and ToM tasks

Finally, we examined whether any brain regions are commonly involved in both reward processing and social cognition. To this end, we tested whether activity in the social cognition-related ROIs was modulated by reward expectation or outcome in the MID task, and conversely, whether activity in the reward-related ROIs was modulated by social cognition during the ToM task.

Several social cognition-related ROIs were found to be associated with reward processing (Fig. 3ab). Neural activity in these ROIs was significantly modulated by the magnitude of monetary reward presented during the Cue phase (*t* = 2.425, *p* = 0.031 in the left TPJ; *t* = 2.674, *p* = 0.019 in the right TPJ; *t* = 2.309, *p* = 0.038 in the left TP; *t* = 2.033, *p* = 0.063 in the right TP; *t* = 2.459, *p* = 0.029 in the dmPFC; *t* = 2.743, *p* = 0.048 in the PCC) and/or by the contrast between success and failure during the Outcome phase of the MID task (*t* = 2.460, *p* = 0.029 in the left TPJ; *t* = 0.507, *p* = 0.621 in the right TPJ; *t* = 2.601, *p* = 0.022 in the left TP; *t* = 1.353, *p* = 0.199 in the right TP; *t* = 2.517, *p* = 0.026 in the dmPFC; *t* = 3.690, *p* = 0.003 in the PCC). However, after applying Bonferroni correction for multiple comparisons across ROIs, only the posterior cingulate cortex (PCC) remained significant, suggesting that the PCC may play a pivotal role in both reward processing and social cognition.

In contrast, none of the reward-related ROIs showed significant modulation by social cognition (Fig. 3c). Specifically, there were no significant differences in neural activity between the Social and Nonsocial conditions of the ToM task within these regions (*t* = −0.434, *p* = 0.672 in the left ventral striatum; *t* = −0.104, *p* = 0.919 in the right ventral striatum; *t* = 1.148, *p* = 0.272 in the vmPFC).

**Figure 3:**
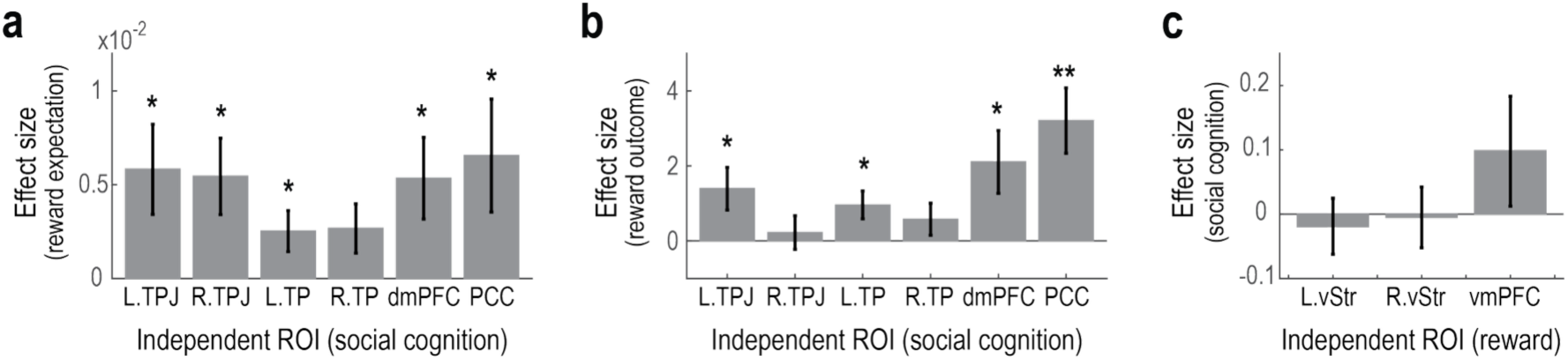
Shared representation between MID and ToM tasks. (a) Neural correlates of reward expectation during the MID task on the social cognition-related ROIs. Bar plots show the effect sizes of reward expectation (i.e., the parametric modulation by the amount of monetary reward presented during the Cue phase) (mean ± SEM across participants). L.TPJ, left temporoparietal junction; R.TPJ, right temporoparietal junction; L.TP, left temporal pole; R.TP, right temporal pole; dmpFC, dorsomedial prefrontal cortex; and PCC, posterior cingulate cortex. \**p* < 0.05. (b) Neural correlates of reward outcome during the MID task on the social cognition-related ROIs. Bar plots show the effect sizes of reward outcome (i.e., the contrast between success and failure during the Outcome phase) (mean ± SEM across participants). \*\**p* < 0.01. (c) Neural correlates of social cognition during the ToM task on the reward-related ROIs. Bar plots show the effect sizes of social cognition (i.e., the contrast between Social and Nonsocial conditions) (mean ± SEM across participants).

## Discussion

This study aimed to replicate previous findings on the neural correlates of reward processing and social cognition in a Japanese sample. To this end, we conducted a functional MRI (fMRI) experiment using the Monetary Incentive Delay (MID) task (Knutson et al. 2000) and the Theory of Mind (ToM) task (Dodell-Feder et al. 2011; Ogawa, Yokoyama, and Kameda 2017) with participants who were native Japanese speakers.

While the MID and ToM tasks are widely used around the world to investigate neural activity related to reward processing and social cognition, prior research has also reported cultural differences in the underlying neural mechanisms (Han and Northoff 2008; Kitayama and Uskul 2011). For example, one study demonstrated that, during the MID task, European American and Chinese participants exhibited differential neural responses to social rewards in the ventral striatum (nucleus accumbens) (Blevins et al. 2023). Furthermore, in a false-belief task—a common Theory of Mind paradigm—differences in inferior frontal gyrus activation were observed between American English-speaking monolinguals and Japanese-English bilinguals (Kobayashi, Glover, and Temple 2006).

In the MID task, we observed that reward expectation and reward outcome were associated with neural activity in the ventral striatum and ventromedial prefrontal cortex (vmPFC), consistent with previous findings (Knutson et al. 2000; Oldham et al. 2018; Jauhar et al. 2021). In the ToM task, social cognition was associated with activation in the temporoparietal junction (TPJ) and temporal pole (TP), aligning with prior studies (Schurz et al. 2014). In contrast, we did not observe significant activation in the dorsomedial prefrontal cortex (dmPFC) associated with social cognition. Overall, these results broadly replicate and extend earlier findings—with the exception of dmPFC activation in the ToM task—and support the cross-cultural generalizability of the neural mechanisms underlying reward processing and social cognition.

Moreover, we found that the posterior cingulate cortex (PCC) was activated in response to both reward processing and social cognition, suggesting a potential integrative role for this region in economic decision-making. This finding aligns with recent theories proposing the PCC as part of a domain-general network for internally guided cognition, serving as a neural bridge between reward evaluation and social inference. The PCC has been implicated in self-referential processing (Northoff et al. 2006), reward-seeking decision-making (Pearson et al. 2011), and mentalizing (Li, Mai, and Liu 2014), and is recognized as a core component of the default mode network (Menon 2023). Together, these findings highlight the PCC as a potential hub integrating reward-related and social cognitive processes.

In conclusion, this study supports the generalizability of previously reported neural and behavioral correlates of reward processing and social cognition by successfully replicating these findings in a Japanese sample. The successful replication also affirms the validity of our fMRI protocols for use in future neuroeconomics research.

## Acknowledgments

This research was supported by the JSPS KAKENHI Grant Number 22K21357 (S.S.).

## Data availability

Behavioral data and code are available at Open Science Framework (https://osf.io/ahv9m/). MRI data are available at OpenNeuro (https://openneuro.org/datasets/ds006401/versions/1.0.0).

---

**Figure S1:**
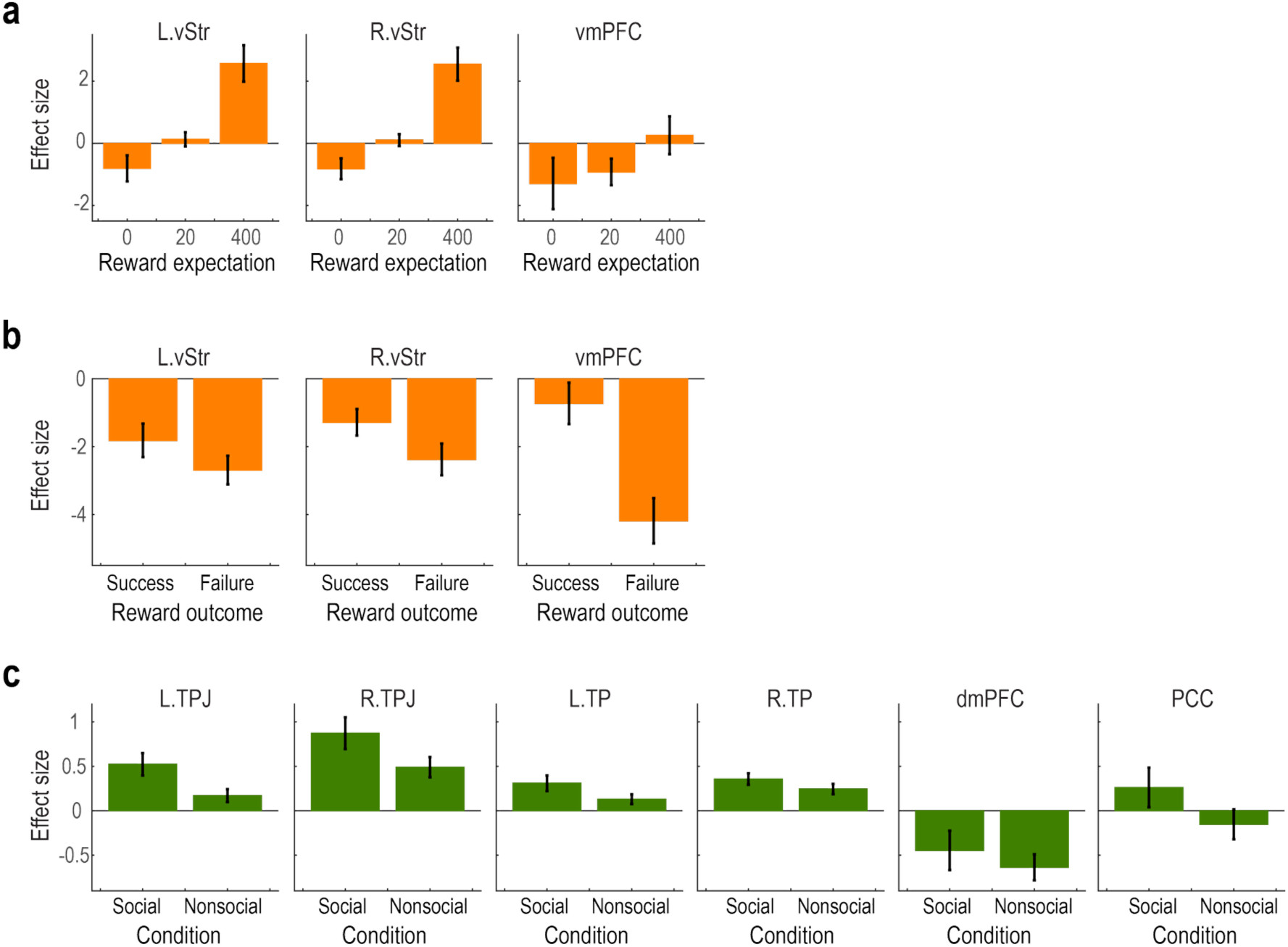
Supplementary analysis on the neuroimaging data. (a) Independent ROI analysis of reward expectation in the MID task. Effect sizes (signal changes from baseline) in the Cue phase are plotted as a function of reward expectation (i.e., the monetary amount available upon a successful outcome) (Mean ± SEM across participants). L.vStr, left ventral striatum; R.vStr, right ventral striatum; and vmPFC, ventromedial prefrontal cortex. (b) Independent ROI analysis of reward outcome in the MID task. Effect sizes (signal changes from baseline) in the Outcome phase are shown separately for success and failure outcomes (Mean ± SEM across participants). (c) Independent ROI analysis of social cognition in the ToM task. Effect sizes (signal changes from baseline) in are shown separately for the Social and Nonsocial conditions (Mean ± SEM across participants). L.TPJ, left temporoparietal junction; R.TPJ, right temporoparietal junction; L.TP, left temporal pole; R.TP, right temporal pole; dmpFC, dorsomedial prefrontal cortex; and PCC, posterior cingulate cortex.

**Table S1.** Brain areas exhibiting significant changes in the BOLD signal associated with reward processing in the Monetary Incentive Delay task.

| Contrast | Region | Hemi | x | y | z | t-statistic | p-value | Voxels |
| --- | --- | --- | --- | --- | --- | --- | --- | --- |
| Reward expectation | Large cluster including ventral striatum | L/R | 30 | -7 | -10 | 7.33 | 0.000 | 9961 |
|  | Temporoparietal junction | L | -54 | -49 | 41 | 6.41 | 0.000 | 228 |
| Reward outcome | Large cluster including dorsal striatum and vmPFC | L/R | 0 | 8 | 14 | 15.71 | 0.000 | 5222 |
|  | Primary motor cortex | L/R | 0 | -28 | 68 | 8.46 | 0.000 | 1137 |
|  | Dorsolateral prefrontal cortex | R | -21 | 38 | 50 | 8.35 | 0.000 | 275 |

Activated clusters observed in the whole-brain analysis ( $P < 0.05$ cluster-level corrected) of fMRI. vmPFC: ventromedial prefrontal cortex. The unthresholded activation maps are available at NeuroVault (https://neurovault.org/images/901831/ and https://neurovault.org/images/901832/).

**Table S2.** Brain areas exhibiting significant changes in the BOLD signal associated with social cognition in the Theory of Mind task.

| Contrast | Region | Hemi | x | y | z | t-statistic | p-value | Voxels |
| --- | --- | --- | --- | --- | --- | --- | --- | --- |
| Social cognition | Temporal pole | R | 54 | 8 | -19 | 6.09 | 0.000 | 232 |
|  | Primary visual cortex | L/R | 6 | -76 | -1 | 5.94 | 0.000 | 260 |
|  | Temporoparietal junction | R | 54 | -46 | 17 | 4.90 | 0.000 | 125 |

Activated clusters observed in the whole-brain analysis ( $P < 0.05$ cluster-level corrected) of fMRI. The unthresholded activation map is available at NeuroVault (https://neurovault.org/images/901812/).

